# APPROACHES TO DIMENSIONALITY REDUCTION FOR ULTRA-HIGH DIMENSIONAL MODELS

**DOI:** 10.1101/2024.08.20.608783

**Authors:** Krzysztof Kotlarz, Dawid Słomian, Joanna Szyda

**Affiliations:** Biostatistics Group, Department of Genetics, Wroclaw University of Environmental and Life Sciences, Wroclaw 51-631, Poland; National Research Institute of Animal Production, Balice 32-083, Poland

**Author notes:** To whom correspondence should be addressed.; **Corresponding authors:** Joanna Szyda.

## Abstract

The rapid advancement of high-throughput sequencing technologies has revolutionised genomic research by providing access to large amounts of genomic data. However, the most important disadvantage of using Whole Genome Sequencing (WGS) data is its statistical nature, the so-called p>>n problem. This study aimed to compare three approaches of feature selection allowing for circumventing the p>>n problem, among which one is a novel modification of Supervised Rank Aggregation (SRA). The use of the three methods was demonstrated in the classification of 1,825 individuals representing the 1000 Bull Genomes Project to 5 breeds, based on 11,915,233 SNP genotypes from WGS. In the first step, we applied three feature (i.e. SNP) selection methods: the mechanistic approach *(SNP tagging)* and two approaches considering biological and statistical contexts by fitting a multiclass logistic regression model followed by either 1-dimensional clustering *(1D-SRA)* or multi-dimensional feature clustering *(MD-SRA)* that was originally proposed in this study. Next, we perform the classification based on a Deep Learning architecture composed of Convolutional Neural Networks. The classification quality of the test data set was expressed by macro F1-Score. The SNPs selected by *SNP tagging* yielded the least satisfactory results (86.87%). Still, this approach offered rapid computing times by focussing only on pairwise LD between SNPs and disregarding the effects of SNP on classification. *1D-SRA* was less suitable for ultra-high-dimensional applications due to computational, memory and storage limitations, however, the SNP set selected by this approach provided the best classification quality (96.81%). *MD-SRA* provided a very good balance between classification quality (95.12%) and computational efficiency (17x lower analysis time and 14x lower data storage), outperforming other methods. Moreover, unlike *SNP tagging*, both SRA-based approaches are universal and not limited to feature selection for genomic data. Our work addresses the urgent need for computational techniques that are both effective and efficient in the analysis and interpretation of large-scale genomic datasets. We offer a model suitable for the classification of ultra-high-dimensional data that implements fusing feature selection and deep learning techniques.

## INTRODUCTION

The past several decades have seen a significant expansion in the availability of genetic data for a rapidly growing number of individuals. Due to the decreasing cost of applying high-throughput technologies, researchers have been able to examine genomes with DNA sequence accuracy [1]. However, this increasing availability of genomic data comes with increasing challenges associated with its storage, bioinformatic processing, and statistical analysis. **Storage** not only requires large hard disk spaces but is also associated with considerable wall clock time spent reading data into programs and writing intermediate and final result files - activities that are very difficult or often impossible to parallelise and execute on GPU. Parallelisation of data processing on CPU (achieved by code vectorisation, applying OpenMP, or even MPI directives) or GPU is typically possible for **bioinformatic data processing** and thus enables faster computations. Still, often limited factors are the number of available parallel processing units and RAM, especially if data cannot be analysed on HPC architectures due to access or privacy constraints. However, the main focus of our study is on **statistical problems** related to high-dimensional genomic data processing [2–4]. They comprise:

i. Problems with accurate model parameter estimation. More specifically, as demonstrated by Giraud [2], standard errors of estimates 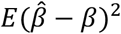linearly increase with the increasing number of model dimensions that, as a consequence, make statistical and biological inferences based on 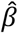inaccurate. Furthermore, when the variation of a dependent variable is described by numerous independent covariates (features), false positive associations may arise due to fitting patterns to “noise” [5]. The accurate estimation of model parameters in a high dimensional setup is also hampered by numerical inaccuracies that may result from many numerical operations, often in the vicinity of over/underflow.
ii. Interpretability of model parameters due to the presence of correlation between subsets of features that varies from high to no correlation, including the thread of fitting false positive patterns of the variation of the dependent variable to noise that arises from the variability of multiple features.
iii. Applicability of traditional hypothesis testing, based on P-values for which the multiple testing correction is often invalid due to violating the underlying assumption of independence between particular tests.
iv. Especially for the specific task of classification, in a multidimensional space, very many data points remain isolated on the boundary of the parameter space, which hampers their assignment to a particular class.

Still, in reality, especially in the case of whole genome sequence data, the underlying true dimensionality of the model is not as complex as the number of available features. Therefore, feature selection (FS) is not only essential to circumvent the challenges mentioned above, but also helps to identify biologically relevant features for downstream analysis. The literature suggests a set of FS techniques widely categorised into basic, hybrid, and ensemble approaches [6]. Our study concentrated on an ensemble approach designed for ultra-high-dimensional data. The ensemble approaches combine several models created from the same dataset to provide the best accuracy and robustness of the feature selection process [7]. To circumvent the above-mentioned problem with P values, the final selection of relevant features is based on a cut-off that is either arbitrarily defined or estimated from the data. An essential step in the ensemble approach to feature selection is rank aggregation (RA), which combines feature importance scores from many models. RA creates an overall rating of features by combining the internal ranking of features within sub-models characterised by different model performances. Feature ranking within a particular sub-model can be based, for example, on feature estimates, while sub-models performance is typically expressed by sub-model fit quality. The final RA, which is often statistically and computationally the most challenging step, can imply a range of aggregation strategies. For instance, a linear mixed model (LMM) can be used due to its robustness towards the violation of asymptotic properties [8] such as normality or homoscedasticity, its ability to handle complex data structures by the possibility of incorporating a feature correlation information directly into the model, as well as by elegant handling of p >> n problem through imposing a *N*(*μ, σ*^2^) shrinkage on model estimates [9].

In our analysis, the problem of data storage was addressed by implementing memory mapping, which allowed us to avoid holding the entire dataset in memory, as well as by CPU- and GPU-based task parallelisation and vectorisation that was directly implemented in rank aggregation procedure and Deep Learning (DL) classification. However, the major goal of our study was to compare three feature selection approaches of varying computational and statistical complexity to define an optimal subset of features for the Deep Learning-based multiclass classification in an ultra-high-dimensional setup of strongly correlated features. For this purpose, a whole genome sequence (WGS) data of 5,063 bulls characterised by 33,595,340 Single Nucleotide Polymorphisms (SNPs) was used to illustrate the three feature selection procedures, for which the goal was to provide classification to five classes represented by breeds. The FSs implement a purely mechanistic approach of reducing the correlation between SNPs expressed by Linkage Disequilibrium (LD), without considering their biological context *(SNP tagging)* and two approaches considering biological and statistical contexts by assessing the importance of SNPs on classification by fitting a multiclass logistic regression model followed up by 1-dimensional clustering *(1D-SRA)* or multi-dimensional feature clustering *(MD-SRA)*.

The first part of the study (i) describes the methodology of SNP selection and breed classification, next (ii) we explore the differences in resulting sets of selected SNPs and the computational efficiency of the applied methods, then (iii) we focus on the quality of multi-class classification via DL-based architectures using the three different SNP sets identified in part (i), while in the last section (iv) we conclude the study with a discussion of major differences between the FS approaches, recommendation of use, and draw directions for future research in this area.

## MATERIAL AND METHODS

### Sequenced individuals and their genomic information

The 1000 Bull Genomes Project Run 9 consists of 5,063 whole genome sequences of *Bos taurus* bulls with 33,595,340 bi-allelic SNP genotypes called by using the GATK software [10] and imputed and phased by Beagle 4.0 [11]. Coverage across these sequences ranged from 1.97x to 173.10x. The dataset incorporates 140 different breeds. The subset of this dataset analysed in our study (Figure 1) comprised individuals with international IDs, characterised by a genome averaged coverage of at least 8, and belonging to breeds represented by at least 150 individuals. SNP preprocessing was done by bcftools 1.10.2 [12] in a parallel mode using 25 threads and involved removing polymorphisms with a call rate lower than 95%, with a minor allele frequency below 5%, as well as multiallelic SNPs. After filtering, 1,825 individuals from five breeds: Angus, Brown Swiss, Holstein, Jersey, and Norwegian Red, were included in the downstream analysis, resulting in 11,915,233 SNPs per individual. This data set was divided into the training dataset of 1,460 individuals and the test dataset of 365 individuals. The split was done in the way to obtain the balanced set of breeds between training and test dataset.

**Figure 1.**
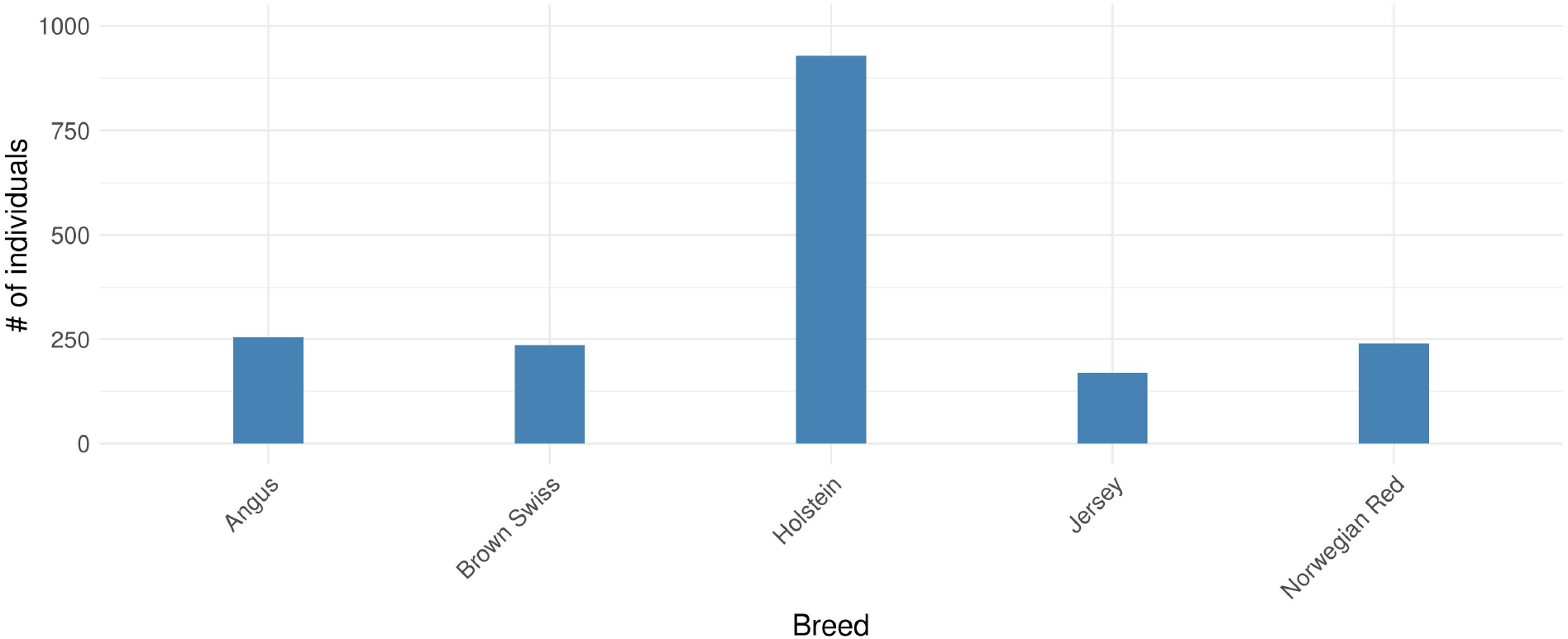
Number of selected individuals across 5 breeds used in the downstream analysis.

### Feature selection

The subset of SNPs and animals defined by the *training data* set was used to select features, i.e. SNPs for the DL-based classifier. The approaches applied for feature selection were described below.

#### SNP aggregation based on linkage disequilibrium (SNP tagging)

This approach was based solely on reducing the correlation between SNPs without considering their biological context. PLINK 1.9 [13] was applied to select SNPs (tagSNPs) representative of a given genomic region based on local Linkage Disequilibrium. The LD between pairs of SNPs was quantified using the R^2^ metric and the tagSNPs were selected for genomic regions spanning 100,000 bp considering a moving window of 1,000 bp. The maximum LD threshold for determining a tag SNP was R^2^ = 0.5.

#### SNP selection-based one-dimensional supervised rank aggregation (1D-SRA)

This approach integrates both biological and statistical contexts by assessing the importance of SNPs for the classification by fitting a multiclass logistic regression model and thus adding the biological component to the feature selection process.

The Supervised Rank Aggregation (SRA) methodology implements supervised learning and rank aggregation for feature selection. The procedure involved the following steps (Figure 2A):

**Figure 2.**
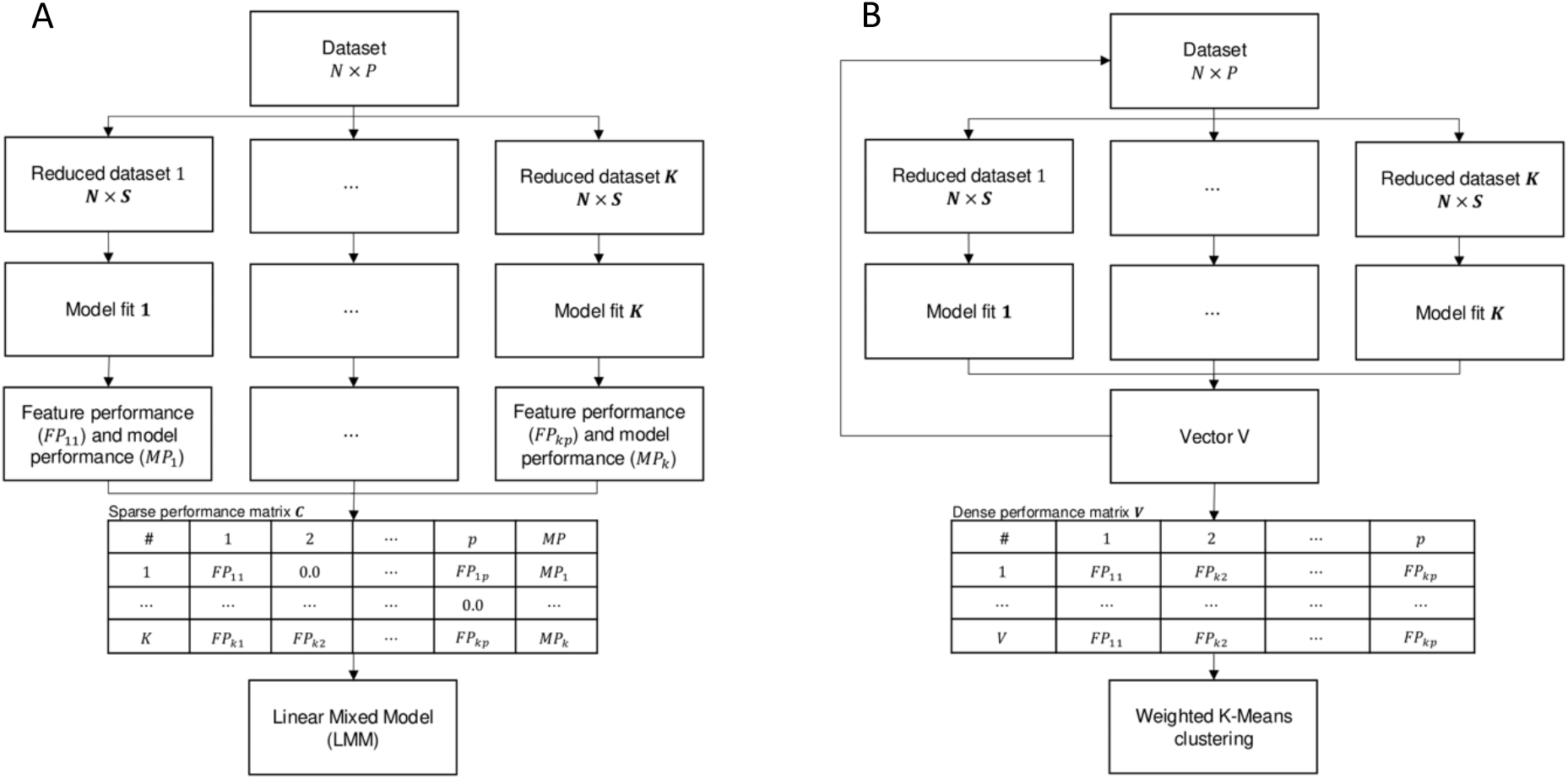
Representation of feature selection procedure implementing A. *1D-SRA* and B. *MD-SRA*.

1. From the dataset of N=1,825 individuals and P=11,915,233 SNPs, K=47,660 reduced data sets composed of all individuals but with randomly selected *S*=250 SNPs were sampled.
2. For each of the reduced data sets, the following K multinomial logistic regression models were fitted for B=5 breed classification (equation 1):

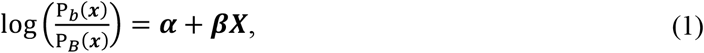

where *P*_*b*_ represents the estimated breed classification probability encoded as an NxB incidence matrix, ***α*** is a 1xB vector of breed-specific intercepts, ***β*** is a BxS matrix of breed-specific SNP estimates with the corresponding design matrix ***X*** containing SNPs genotypes coded as: 0/0 represented as 0, 0/1 represented as 1 and 1/1 represented as 2.
3. Model Performance of each of the K models (*MP*_*k*_) was expressed by the quality of fit of each of the multinomial logistic regression models quantified by the cross-entropy loss (CE) metric (equation 2):

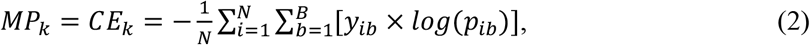

where *y*_*ib*_ represents the true classification of individual *i* to breed *b* and *p*_*ib*_ is the corresponding probability estimated by model (1) that is given either by 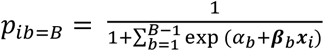 for the reference breed (*B*) or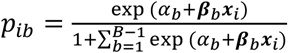 for each non-reference breed *b*.
4. The resulting model performance matrix ***C*** is a sparse matrix of P columns containing features’ performances (FPs) and *MP* in the *P* + 1 column, and K rows corresponding to each reduced model. Precisely, each row contains FPs of S SNPs fitted in *k*-th reduced model, expressed by *max*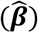 for each SNP, while FPs of the remaining P-S SNPs were set to zero, and the quality of fit of this model was expressed by 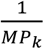 Due to the large dimension of ***C***, its disk storage was resolved by using the memory map approach implemented in Numpy 1.24.3 [14].
5. The aggregation step comprised combining effects of all SNPs from multinomial logistic regression reduced models into a single mixed linear model (LMM), given by:

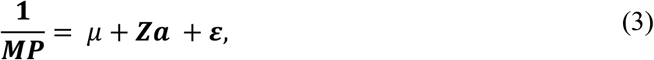

where **MP** [Kx1] represents the vector of fit of multiclass regression reduced models expressed by the reciprocal of cross-entropy loss ^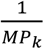^, *μ* is a general mean, ***a*** [Px1] is a vector of random SNP effects with a pre-imposed normal distribution *N*(0, ***I****δ*_*a*_^2^), with the corresponding design matrix ***Z*** [KxP] containing *max*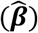 of SNP effects resulting from the reduced models and is thus equivalent to the first P columns of the model performance matrix ***C***, and **ε** [Kx1] is a vector of random residuals distributed as *N*(0, *Iδ*_*ε*_^2^). The model parameters were estimated by the mix99 software [15], assuming the variances of 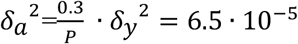 and 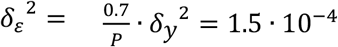 as known (i.e. not estimated) since in such a large system of dense equations *δa*^2^ and *δ*_*ε*_^2^ would have been technically impossible to estimate. Moreover, the goal of fitting the model is not to estimate SNP effects but to provide a ranking of features that will further be used for DL-based classification. Due to the large dimensions of the equation system, a direct solution by solving the mixed model equations was not possible. Therefore, the iterative preconditioned conjugate gradient (PCG) method was applied, in which the preconditioned matrix was defined based on block diagonal elements of the coefficient matrix from mixed model equations [16]. The CD convergence criterion of 1.0. 10^−3^ was set as the stopping criterion for PCG and the solver was run in parallel on 40 CPUs.
6. The final step in the feature selection process comprised the grouping of the SNP effect estimates 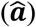 into sets *relevant* and *irrelevant* for classification, so that the relevant features will further be used to train the DL-based classifier. This was done by applying the 1-dimensional K-means classification algorithm for defining two clusters, implemented in the Scikit-learn 1.3.0 library [17]. Using K-means allowed for avoiding setting up an arbitrary cutoff value to define the *relevant* and *irrelevant* SNP sets. This procedure assumed that features with greater importance for classification resulted in models with high MP, which corresponds to low cross-entropy loss [6].

#### SNP selection-based multidimensional-dimensional rank aggregation (MD-SRA)

Due to the very high dimensions of the equation system underlying LMM, which was used for feature aggregation in *1D-SRA* and was demanding computationally, the alternative, less computationally heavy approach was proposed. In this approach, the aggregation of features by LMM was substituted by the direct clustering of multiple model performance matrices ***C*** aggregated on all reduced models with multidimensional K-means. The procedure comprised the following steps (Figure 2B):

1. Five model performance matrices ***C*1-*C*5** were generated by repeating the steps 1-4 described above. Since the performance of a single feature occurs only once within each model performance matrix, K rows of each model performance matrix were collapsed to form one dense vector of non-zero FPs, forming ***V*1-*V*5** model performance vectors.
2. *Relevant* and *irrelevant* groups of features were defined by assuming two clusters in five-dimensional weighted K-means clustering of matrix ***V*** that was formed by appending dense vectors ***V*1-*V*5**. The K-means algorithm was weighted by the multiplication of each element(*FP*_*ij*_)of ***V*** by the reciprocal of cross-entropy loss 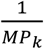 of the corresponding the reduced model. The L-2 norm of the five-dimensional coordinates of the cluster centroids was used to define the cluster, containing the *relevant* features, which corresponds to the distance of each centroid in the five-dimensional space.

### Classification

The classification was based on the DL and was implemented in Python via the Keras interface [18] to the TensorFlow library [19]. The following architecture was implemented (Figure 3):

**Figure 3.**
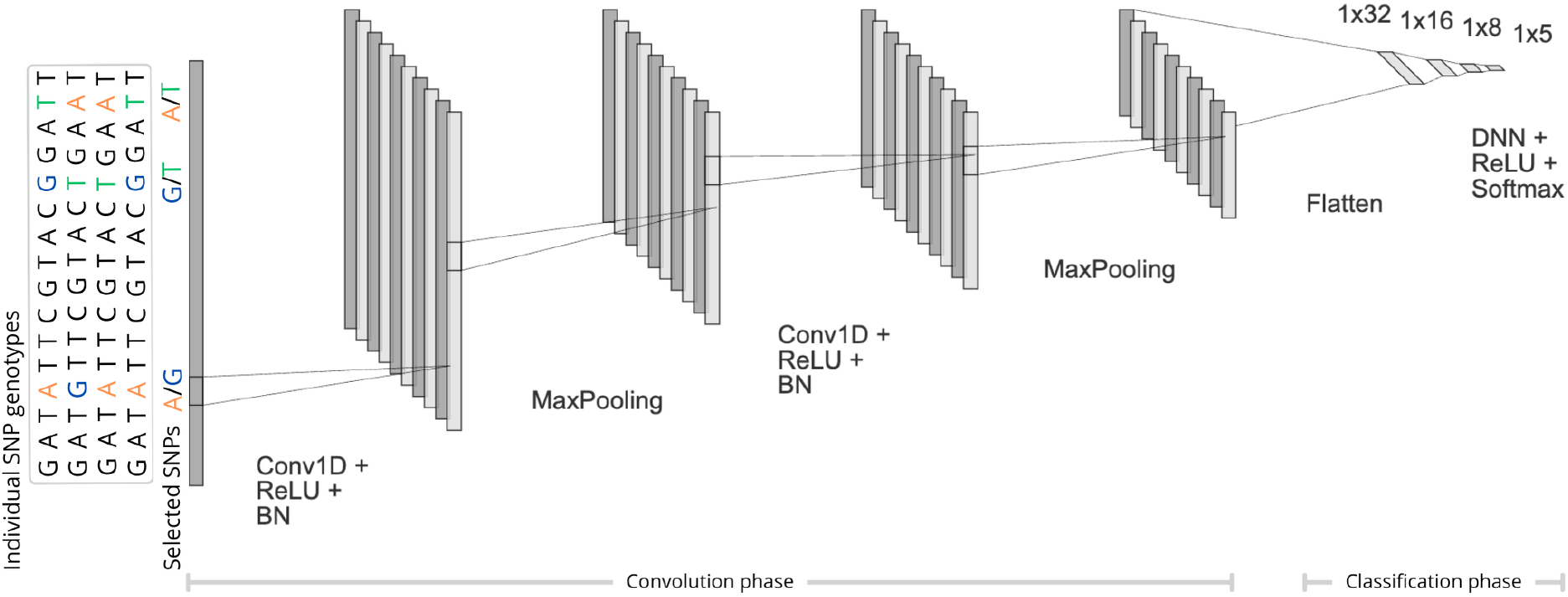
The architecture of the Deep Learning-based classifier. The individual genotypes of SNPs originate from three feature selection methods: *SNP tagging, 1D-SRA*, or *MD-SRA*.

1. The input consisted of the SNP genotypes that resulted from each of the three feature selection approaches (*SNP tagging, 1D-SRA*, and *MD-SRA*) described above and of the breed class assignments for each individual (five breeds).
2. First, in the convolution phase, (i) a sequential 1D Convolutional Neural Network (1D-CNN) was applied, followed by the Rectified Linear Unit (ReLU) activation function *f*_*ReLU*_(*Z*_*i*_) = *max*(0, *Z*_*i*_), (ii) a batch normalization (BN) layer, and (iii) a max pooling layer with pool size equal to five. In each 1D convolutional layer, 10 kernels of size 150 were applied to the input to capture local patterns by sliding across the input. After each 1D-CNN, a batch normalization layer was used to normalise the output, so that each layer received input with the same mean and variance. This was followed by a max-pooling layer, which selected the maximum value from each feature map window to decrease the dimensionality (Figure 3).
3. In the last section of the DL architecture, the output of 1D-CNN was flattened to a 1-dimensional vector processed by three hidden dense layers with the ReLU activation function (Figure 3).
4. The final classification was performed by the last DNN implementing the softmax activation function: 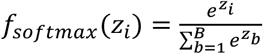 where *Z*_*i*_ represents the output of the last layer and *B* is the number of classes (i.e. breeds).

The Adam optimiser [20] implementing the stochastic gradient descent algorithm with a default learning rate of 5.0. 10^−5^ was applied to minimise the categorical focal-cross-entropy loss (*FL*) function given by:

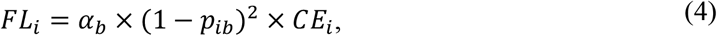

where 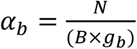 was used as suggested by Li et al. [21], with *N* representing the number of individuals, *B* = *5* representing the number of classes, and *g*_*b*_ being the number of individuals assigned to class *b* in the training dataset. FL was implemented to mitigate the class frequency imbalance during optimisation, by penalising the impact of more numerous.

### DL-based classifier evaluation

To evaluate the classification performance of the DL-based classifier during the training phase, stratified 5-fold cross-validation was applied by randomly selecting 80% of the training individuals for training the model and leaving out the remaining 20% of the individuals for the validation dataset. The test dataset was used to assess the final performance of the trained model. The quality of classification was quantified using the Macro F-1 Score metric (equation 5):

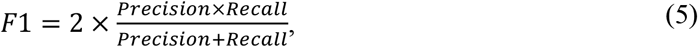

where 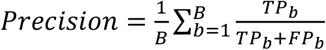 and 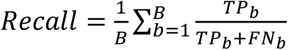 the *TP*_*b*_ (true positive) class representing the individuals correctly classified to breed *b*, the *FN*_*b*_ (false negative) class representing the individuals of breed b incorrectly classified to another breed, the *FP*_*b*_ (false positive) class representing the individuals from another breed incorrectly classified as breed b. Additionally, the AUC (Area Under the Curve) resulting from averaging AUCs for each individual corresponding to ROC (Receiver Operating Characteristic) was constructed separately for the classification of each breed. The models were trained and evaluated on the NVIDIA Tesla P100 GPU with 328 GB of RAM.

## RESULTS

### Feature selection

In this section, we present the results of SNP pre-selection by using the three proposed dimensionality reduction methods: (i) *SNP tagging* based on LD, (ii) *1D-SRA* based on a linear mixed model, and (iii) *MD-SRA* based on weighted multi-dimensional clustering. The computational efficiency of the SNP selection approaches varied considerably, reflecting the complexity of analysis associated with each approach (Table 1).

**Table 1.**
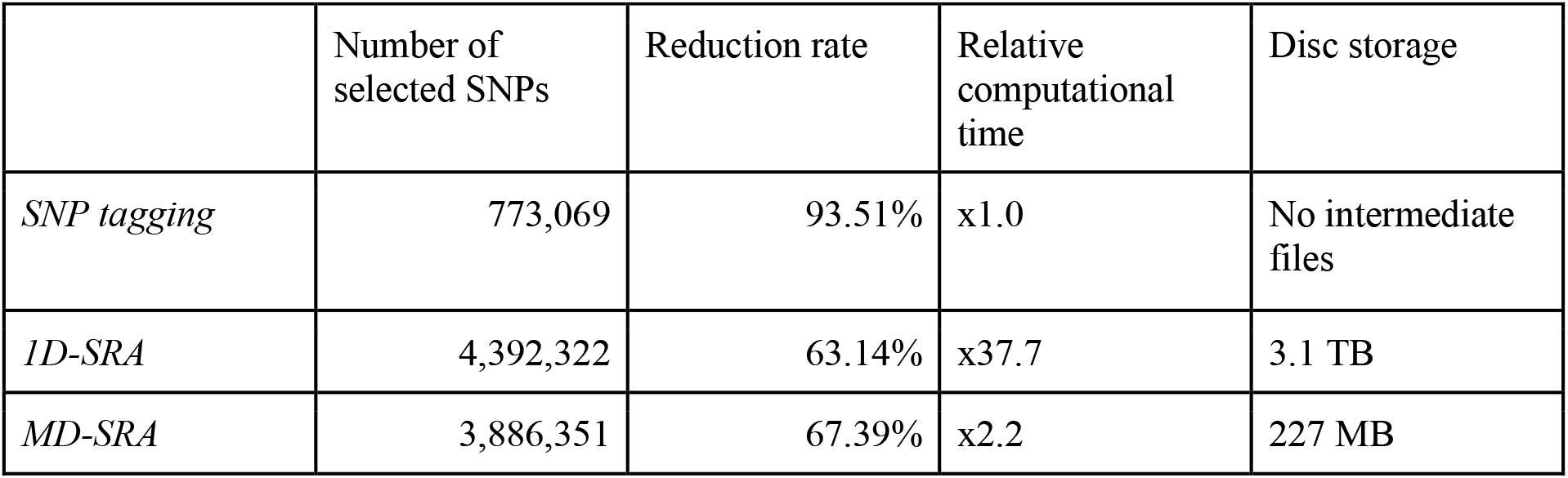
Summary results of the three feature selection approaches.

### SNP tagging

The LD pruning method, being the simplest in terms of computational demand, was set as the baseline for our comparison. It required 74 minutes to complete the SNP selection process. The initial set of 11,915,233 SNPs was reduced to 773,069 SNPs (reduction rate 93.51%). The disk storage requirements were very small since this approach retains only the pruned set of SNPs without intermediate files.

### 1D-SRA

The approach involved fitting multiple multiclass logistic regression models followed by rank aggregation based on LMM. Since multiple logistic regression-reduced models were fitted, this approach required considerable CPU resources. Furthermore, CPU time was needed to compute a design matrix **Z** for LMM and obtain LMM solutions. In addition, the procedure required storing and reading multiple files on a hard disk, without the possibility of completing all *1D-SRA* pipeline in memory. The wall clock time of 2,790 minutes (46 hours and 30 minutes) was 37.7 times longer than for LD pruning. The initial set of SNPs was reduced to 4,392,322 SNPs, achieving a reduction rate of 63.14%. Since this approach required substantial hard disk storage of estimates of the logistic regression model effects needed for the aggregation step by the LMM, it resulted in terabytes of file size (3.1 TB for the dataset analysed in this study), moreover, the Z matrix was also stored on a disk by implementing memory mapping approach and then rewritten into the plain text file, required by the mix99 software.

### MD-SRA

The approach provided the advantage of incorporating statistical benefits and retaining computational efficiency. The wall clock time of 2 hours and 40 minutes was only 2.2 times longer than the LD-based approach. The original set of SNPs was reduced to 3,886,351 SNPs, providing a reduction rate of 67.39%. The hard disk space required for the reduced feature performance matrix (227 MB for the dataset analysed in this study) was markedly lower than in the *1D-SRA* method due to fewer intermediate steps.

Both the *1D-SRA* and *MD-SRA* approaches selected a very similar number of SNPs marking features relevant for classification. However, the overlap in SNPs between both approaches was very low. Interestingly, the *SNP tagging* approach shared an equivalent overlap portion with both *1D-SRA* and *MD-SRA* (Figure 4).

**Figure 4.**
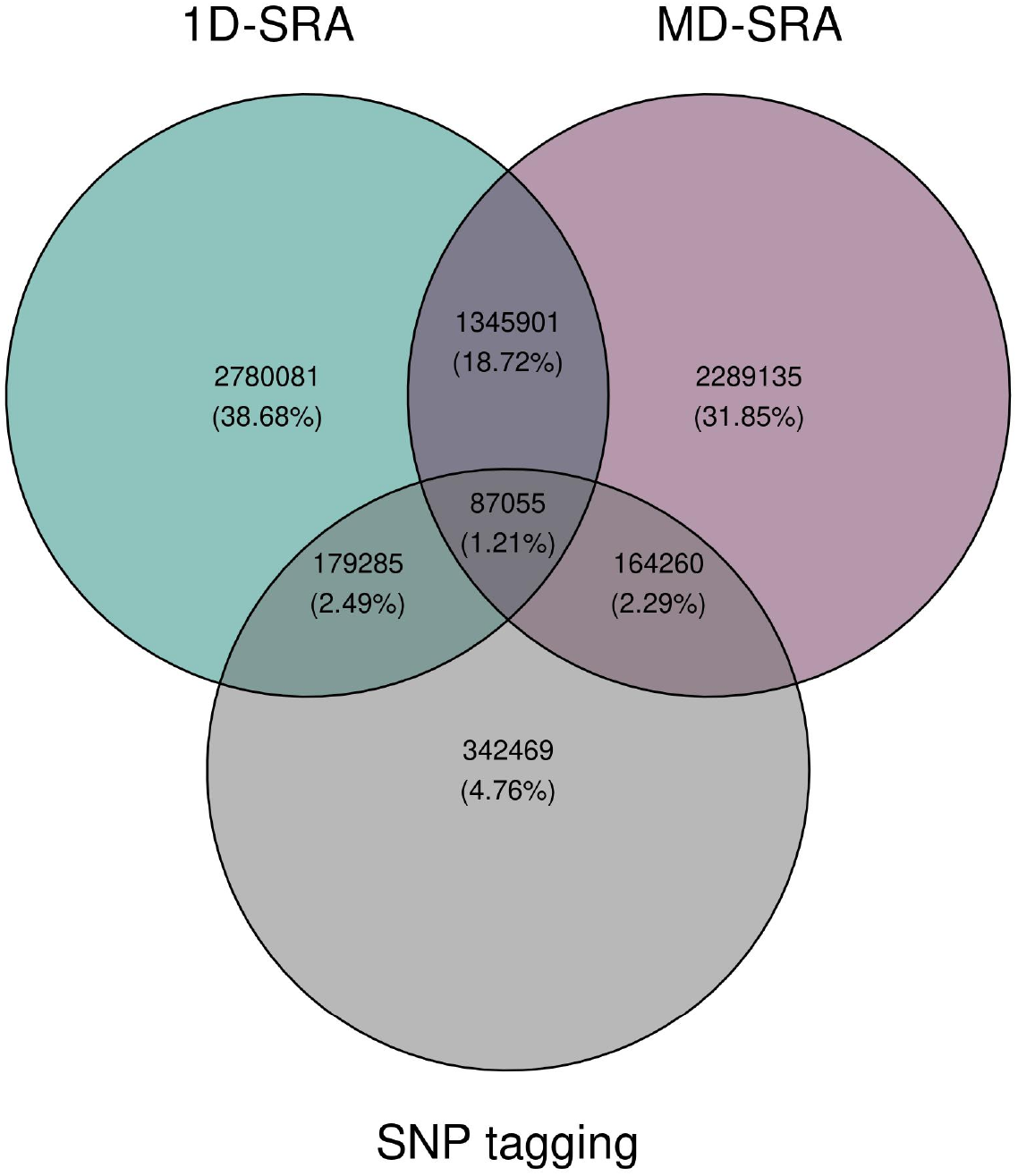
The overlap of relevant SNPs selected by the three feature selection approaches.

### Classification

As summarised in Table 2, all three feature selection approaches resulted in high F1-Scores that indicated a good balance between precision and recall, and the AUC suggested a good ability to separate classes. For the *validation data* sets, the largest AUC (averaged across validation data sets) of 0.992 was achieved for *1D-SRA*, closely followed by *MD-SRA* with the AUC=0.984. *SNP tagging* resulted in the lowest, albeit still high AUC of 0.963. The same ranking of feature selection approaches was obtained, considering the F1-Score metric with *1-D SRA, MD-SRA*, and *SNP tagging* scoring 90.09%, 88.22%, and 74.65% respectively. Apart from a very good classification performance of all three approaches on the validation data sets, considerable differences in their robustness expressed by a standard deviation of the metrics, were observed. *1D-SRA* was the most robust, with the lowest standard deviations of the AUC and the F1-Score. *MD-SRA* was intermediate, with a standard deviation of AUC approximately three times higher and a standard deviation of the F1-Score twice as high as *1D-SRA*. The validation quality of *SNP tagging* was the least robust, with a standard deviation of the AUC being six times higher and a standard deviation of the F1-Score - three times higher than for *1D-SRA*. Regarding the classification quality of the *test data* set expressed by the F1-Score, a very similar situation as for the validation test was observed. *1D-SRA* and *MD-SRA* provided a very similar classification quality of over 95%, while *SNP tagging* resulted in a good, but lower, F1-Score of 86.87%. The AUC was very similar across all three approaches, with *1D-SRA* and *MD-SRA* achieving AUCs of 0.997 and 0.998 respectively, while *SNP tagging* resulted in a slightly lower AUC of 0.985. The confusion matrix for the test dataset across the individuals was shown on Figure 5.

**Table 2.**
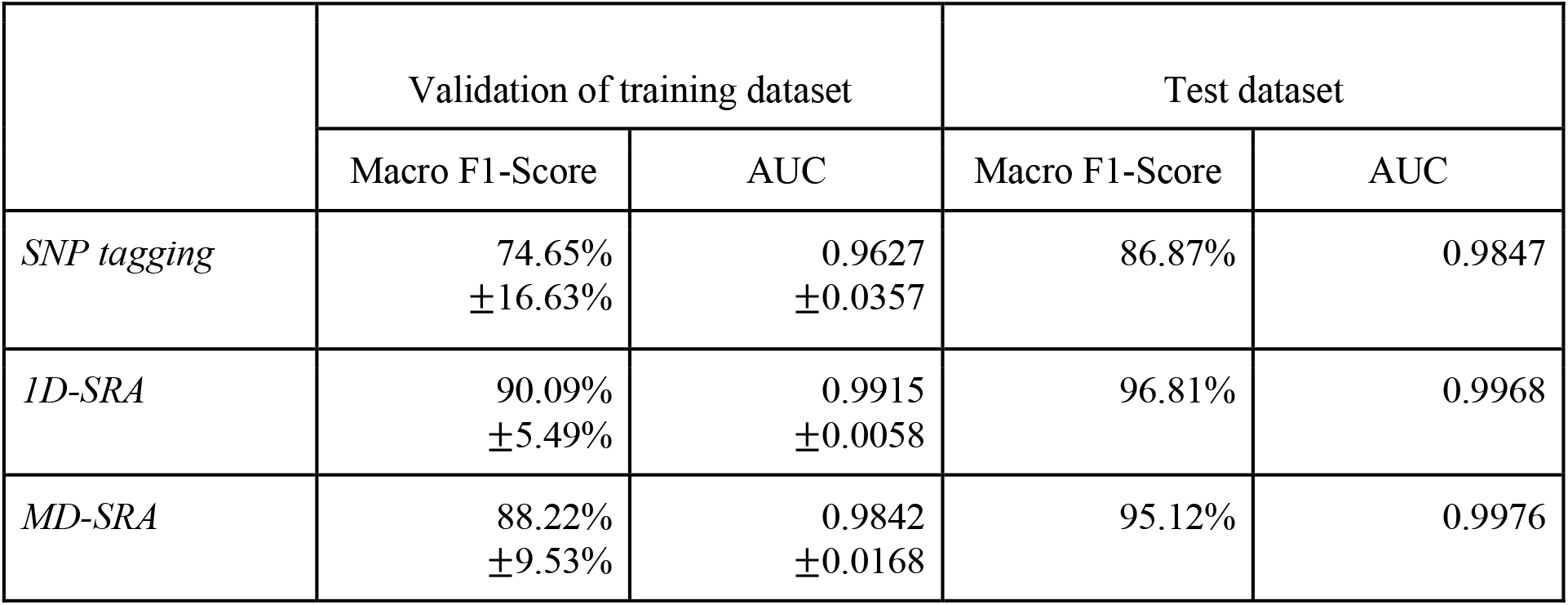
Macro F1-Score and AUC for validation of training datasets.

**Figure 5.**
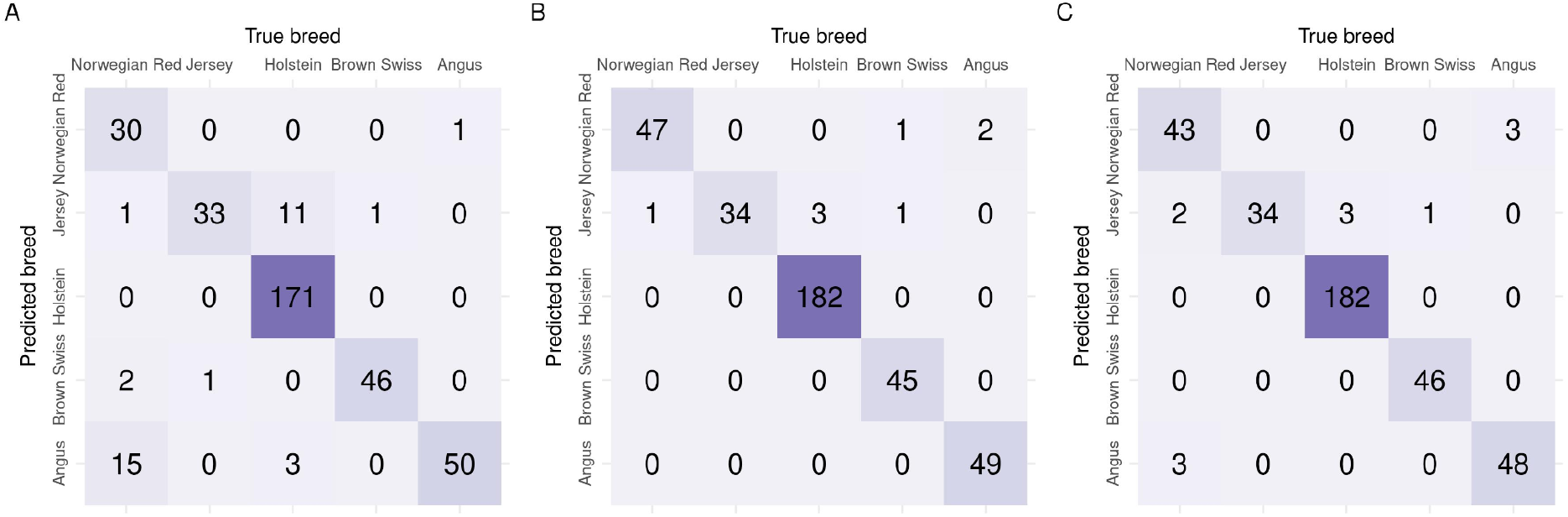
Confusion matrixes of predictions for the test dataset across three feature selection methods A. *SNP tagging* B. *1D-SRA* and C. *MD-SRA*.

## DISCUSSION

Our study explored three procedures of feature pre-selection in ultra-high-dimensional data in the context of classification, illustrated by the classification of bulls into five breeds based on their SNP genotypes identified from WGS. The procedures ranged from a mechanistic approach based solely on exploring pairwise LD between SNPs to computationally and statistically complex approaches incorporating biological information. The proposed methods consider different sources of information contained in a dataset, each with its strengths and limitations. *SNP tagging* is a well-known approach highly used in genomics since it was explicitly designed to prune polymorphic genetic variants, such as SNPs, before further downstream analyses [22]. Since this method is limited only to this specific data structure (SNPs), its high computing efficiency makes it a reasonable option for feature selection, albeit only in genomics. The obvious disadvantage is that it cannot be applied to other types of highly dimensional data. *1D-SRA* is a general approach [6] it has a wide range of applications that extend beyond biology. However, this increased flexibility comes with a higher data processing cost and computational complexity that increases with increasing dimensions of the datasets. Due to its complexity, maximal handlebar dimensions are restricted by available computational resources, including not only memory but also hard disk space, time, and software. The latter strongly depends on the choice of the aggregation model. In the original application [6] an aggregation via penalised regression or random forest was applied, while we implemented LMM. The advantages of using LMM lay not only in its robustness towards the violation of theoretical model assumptions, e.g. of data normality [23], but mostly in its ability to handle correlated data with a complex covariance structure (although the latter was not required in our study). The ability of LMMs to include both, the feature and the residual covariance directly into the model prevents false positive associations that, in the genomic context, may arise from the ascertainment bias due to population structure or varying degrees of family relationship [24]. *MD-SRA* is a modified approach of Jain and Xu [6] that was proposed *de novo* in our study to circumvent the problems associated with the aggregation model. It aims to balance the advantages of the first two approaches, as it is applicable to high-dimensional datasets not only from the area of biology, but at the same time allows for efficient computing performance and low computing resources usage. It allows for capturing interactions between features without the need of fitting the final aggregation model, which is the most computationally intensive element of *1D-SRA*, but still achieves very similar classification quality, but with significantly reduced computation times as compared to *1D-SRA*. In our application, *MD-SRA* demonstrated 17 times shorter analysis time and 14 times shorter data storage. Due to its good classification performance coupled with high computational efficiency, *MD-SRA* is a desirable feature selection algorithm for ultra-high-dimensional data.

Nowadays, in machine learning, with the advance of large data sets with millions of records and millions of explanatory variables, feature pre-selection aims not only to identify relevant features but also to reduce noise that poses the danger of fitting fake patterns to the data that is not functionally relevant and that impedes prediction quality of new data, non-seen by the trained model. Furthermore, by providing fast-growing sizes of the available data sets, it is crucial to build computationally efficient algorithms [25]. Conventional machine learning methods, such as penalized regression (e.g. LASSO), offer simultaneous feature preselection and feature importance estimation by selecting features that significantly contribute to prediction or classification [6]. However, these methods may not always capture the interactions between features present in high-dimensional data. Moreover, in high dimensional data, feature selection that is solely based on statistical significance expressed by P-values, may no longer be relevant, since the existing multiple testing correction approaches neither scale with the very high multiple testing dimensionality, nor consider the correlation between tests (e.g. the Bonferroni correction) [26]. Therefore, the grouping of features as *relevant* and *irrelevant* for downstream analysis is the currently widely used approach [27]. Also, our study adopts the concept of using the dynamic threshold to differentiate between relevant and irrelevant features. This approach was demonstrated by Jain and Xu [6] who implemented it as the 1D K-means clustering and later extended to multidimensional K-means clustering, enabling the classifier to incorporate various model quality metrics. The algorithm proposed in our study is capable of identifying and prioritizing features by dynamically adjusting the relevant feature selection threshold in a multidimensional space defined by feature estimates from multiple reduced models. Such a low complexity approach as K-means clustering performs accurately and efficiently in multiple dimensions, especially that in multiple dimensions, the identification of clusters corresponding to the relevant features is not straightforward [28].

A significant challenge in feature selection, as highlighted in our study and many others [29,30], is that the ranking of features is based on model fit and not on model prediction quality, so that, feature selection prioritises features that make the model fit the training data well. However, this does not always lead to better predictions of new data. Also, in our study *1D-SRA* and *MD-SRA* used model fit (expressed as *MP*_*k*_) for ranking/weighting of features, while for SNP tagging model fit was not considered. But when the classification performance of the CNN classifier based on features preselected by the three approaches was evaluated, the *1D-SRA* was highlighted as the best method in terms of classification quality and robustness. It demonstrated the highest Macro F1-Score and its lowest variation across cross-validation datasets and also the best class prediction performance for the test dataset. Very similar, only slightly lower, classification metrics were obtained by *MD-SRA*. While high demands of *1D-SRA* on computational resources, limit its application for feature selection in ultra-high-dimensional setup. These, favour the former approach for the practical application of feature selection in the case of ultra-high-dimensional data. SNP tagging produced the least satisfactory results, but still resulted in good performance.

Since, in the advent of ever-growing dimensions of data sets, feature selection is and will be an important data analysis step. Immediate future research application regarding the approach proposed in our study would be the exploration of ways to differentiate between important and unimportant features after ranking as well as the evaluation of the performance of the three approaches to predict quantitative outcomes. Specifically, even though K-means clustering worked well in our investigation, it’s worthwhile to investigate more complex feature clustering techniques like hierarchical clustering, Gaussian Mixture Models, or other advanced methods available. However, these approaches come with disadvantages, especially regarding computational limitations. Advanced clustering methods often require significantly more computational resources, especially when dealing with large datasets leading to longer processing times which may not be feasible in real-time applications [31,32]. It is worth considering that some of the clustering approaches perform better in one-dimensional data than in multi-dimensional spaces [33]. Regarding prediction, it is important (i) to differentiate between the quality of prediction for new data ascertained from a similar or from a different general population, to assess which approach is more robust towards potential ascertainment bias and (ii) to explore the prediction quality for data that are dynamic across an additional dimension, such as e.g. populations undergoing strong natural or artificial selection pressures across time.

## CONCLUSIONS

With the very fast-growing dimensionality of data sets, which comprises not only an increasing number of data records but also an increasing number of features the storage and computational burden is an instantly increasing challenge [3,4]. There is a clear need for effective and efficient computing [34]. *MD-SRA* provides a very good balance between classification quality, computational intensity, and required hardware resources among the three compared methodologies. It demonstrated classification performance very similar to *1D-SRA* and markedly better than SNP tagging. Moreover, in contrast to SNP tagging, it is a universal approach, not constrained to the genomic data.

## DATA AVAILABILITY

VCF files are freely available for the members of the 1000 Bull Genomes initiative or upon request to Amanda J Chamberlain.

## SUPPLEMENTARY DATA

### AUTHOR CONTRIBUTIONS

Krzysztof Kotlarz: Conceptualization, Data curation, Formal analysis, Methodology, Validation, Writing – original draft. Dawid Słomian: Data curation, Formal Analysis, Writing—review & editing. Joanna Szyda: Funding acquisition, Investigation, Project administration, Supervision, Writing – original draft.

## ACKNOWLEDGEMENTS

Computations were carried out at the Wroclaw Centre for Networking and Supercomputing and Poznan Supercomputing and Networking Center.

## FUNDING

This work was supported by the National Science Centre [2019/35/O/NZ9/00237].

## CONFLICT OF INTEREST

None

## Notes

### Competing Interest Statement

The authors have declared no competing interest.

## REFERENCES

1. Routhier E, Mozziconacci J. Genomics enters the deep learning era. PeerJ 2022; 10:e13613

2. Giraud C. Introduction to High-Dimensional Statistics. 2021;

3. Johnstone IM, Titterington DM. Statistical challenges of high-dimensional data. Philosophical Transactions of the Royal Society A: Mathematical, Physical and Engineering Sciences 2009; 367:4237–4253

4. Fan J, Li R. Statistical Challenges with High Dimensionality: Feature Selection in Knowledge Discovery. 2006;

5. Hancer E, Xue B, Zhang M. A survey on feature selection approaches for clustering. Artif Intell Rev 2020; 53:4519–4545

6. Jain R, Xu W. Supervised Rank Aggregation (SRA): A Novel Rank Aggregation Approach for Ensemble-based Feature Selection. Recent Advances in Computer Science and Communications 2024; 17:

7. Viharos ZJ, Kis KB, Fodor Á, et al. Adaptive, Hybrid Feature Selection (AHFS). Pattern Recognit 2021; 116:107932

8. Jiang J. ASYMPTOTIC PROPERTIES OF THE EMPIRICAL BLUP AND BLUE IN MIXED LINEAR MODELS. Stat Sin 1998; 8:861–885

9. Henderson CR. Applications of Linear Models in Animal Breeding. 1984;

10. der Auwera GA, O’Connor BD. Genomics in the cloud: using Docker, GATK, and WDL in Terra. 2020;

11. Browning BL, Zhou Y, Browning SR. A One-Penny Imputed Genome from Next-Generation Reference Panels. The American Journal of Human Genetics 2018; 103:338–348

12. Danecek P, Bonfield JK, Liddle J, et al. Twelve years of SAMtools and BCFtools. Gigascience 2021; 10:

13. Purcell S, Chang C. PLINK 1.9. Available from: https://www.cog-genomics.org/plink/1.9 2015;

14. Harris CR, Millman KJ, van der Walt SJ, et al. Array programming with NumPy. Nature 2020; 585:357–362

15. Martin Lidauer Kmemtpmtis. Technical reference guide for MiX99 solver. 2022;

16. Strandén I, Lidauer M. Solving Large Mixed Linear Models Using Preconditioned Conjugate Gradient Iteration. J Dairy Sci 1999; 82:2779–2787

17. Pedregosa F, Varoquaux G, Gramfort A, et al. Scikit-learn: Machine Learning in Python. Journal of Machine Learning Research 2011; 12:2825–2830

18. Chollet F, others. Keras. 2015;

19. Abadi M, Barham P, Chen J, et al. TensorFlow: A system for large-scale machine learning. 12th USENIX Symposium on Operating Systems Design and Implementation (OSDI 16) 2016; 265–283

20. Kingma DP, Ba J. Adam: A Method for Stochastic Optimization. 2014;

21. Lin T-Y, Goyal P, Girshick R, et al. Focal Loss for Dense Object Detection. IEEE Trans Pattern Anal Mach Intell 2020; 42:318–327

22. Ding K, Kullo IJ. Methods for the selection of tagging SNPs: a comparison of tagging efficiency and performance. European Journal of Human Genetics 2007; 15:228–236

23. Schielzeth H, Dingemanse NJ, Nakagawa S, et al. Robustness of linear mixed-effects models to violations of distributional assumptions. Methods Ecol Evol 2020; 11:1141–1152

24. Yang J, Zaitlen NA, Goddard ME, et al. Advantages and pitfalls in the application of mixed-model association methods. Nat Genet 2014; 46:100–106

25. Adadi A. A survey on data-efficient algorithms in big data era. J Big Data 2021; 8:24

26. Yu L, Liu H. Feature Selection for High-Dimensional Data: A Fast Correlation-Based Filter Solution. Proceedings, Twentieth International Conference on Machine Learning 2003; 2:856–863

27. Kamalov F, Sulieman H, Moussa S, et al. Nested ensemble selection: An effective hybrid feature selection method. Heliyon 2023; 9:e19686

28. Ikotun AM, Ezugwu AE, Abualigah L, et al. K-means clustering algorithms: A comprehensive review, variants analysis, and advances in the era of big data. Inf Sci (N Y) 2023; 622:178–210

29. Barrera-García J, Cisternas-Caneo F, Crawford B, et al. Feature Selection Problem and Metaheuristics: A Systematic Literature Review about Its Formulation, Evaluation and Applications. Biomimetics 2023; 9:9

30. Mamdouh Farghaly H, Abd El-Hafeez T. A high-quality feature selection method based on frequent and correlated items for text classification. Soft comput 2023; 27:11259–11274

31. Pinto RC, Engel PM. A Fast Incremental Gaussian Mixture Model. PLoS One 2015; 10:e0139931

32. Wan H, Wang H, Scotney B, et al. A Novel Gaussian Mixture Model for Classification. 2019 IEEE International Conference on Systems, Man and Cybernetics (SMC) 2019; 3298–3303

33. Zhao Y, Shrivastava AK, Tsui KL. Regularized Gaussian Mixture Model for High-Dimensional Clustering. IEEE Trans Cybern 2019; 49:3677–3688

34. König IR, Auerbach J, Gola D, et al. Machine learning and data mining in complex genomic data— a review on the lessons learned in Genetic Analysis Workshop 19. BMC Genet 2016; 17:S1

